# Genetic variation influences pluripotent ground state stability in mouse embryonic stem cells through a hierarchy of molecular phenotypes

**DOI:** 10.1101/552059

**Authors:** Daniel A. Skelly, Anne Czechanski, Candice Byers, Selcan Aydin, Catrina Spruce, Chris Olivier, Kwangbom Choi, Daniel M. Gatti, Narayanan Raghupathy, Alexander Stanton, Matthew Vincent, Stephanie Dion, Ian Greenstein, Matthew Pankratz, Devin K. Porter, Whitney Martin, Wenning Qin, Alison H. Harrill, Ted Choi, Gary A. Churchill, Steven C. Munger, Christopher L. Baker, Laura G. Reinholdt

**Affiliations:** The Jackson Laboratory, Bar Harbor, Maine 04609; Sackler School of Graduate Biomedical Sciences, Tufts University, Boston, MA 02111; Predictive Biology, Inc., Carlsbad, CA 92010; Division of the National Toxicology Program, the National Institute of Environmental Health Sciences, Research Triangle Park, NC 27709

## Abstract

Mouse embryonic stem cells (mESCs) cultured under controlled conditions occupy a stable ground state where pluripotency-associated transcriptional and epigenetic circuitry are highly active. However, mESCs from some genetic backgrounds exhibit metastability, where ground state pluripotency is lost in the absence of ERK1/2 and GSK3 inhibition. We dissected the genetic basis of metastability by profiling gene expression and chromatin accessibility in 185 genetically heterogeneous mESCs. We mapped thousands of loci affecting chromatin accessibility and/or transcript abundance, including eleven instances where distant QTL co-localized in clusters. For one cluster we identified *Lifr* transcript abundance as the causal intermediate regulating 122 distant genes enriched for roles in maintenance of pluripotency. Joint mediation analysis implicated a single enhancer variant ~10kb upstream of *Lifr* that alters chromatin accessibility and precipitates a cascade of molecular events affecting maintenance of pluripotency. We validated this hypothesis using reciprocal allele swaps, revealing mechanistic details underlying variability in ground state metastability in mESCs.

## Introduction

Derivation and *in vitro* propagation of pluripotent mouse embryonic stem cells (mESCs) is heavily influenced by genetic background. Successful derivation of pluripotent mESC lines was first reported in 1981 using both inbred (substrains of 129, [C3HxC57BL/6] F1 hybrids) and certain outbred laboratory strains (Evans and Kaufman, 1981; Martin, 1981). These early reports demonstrated that, in the presence of serum and exogenous leukemia inhibitory factor (LIF), mESCs are capable of robust propagation *in vitro* and sustained capacity to contribute to all embryonic lineages, including the germ line (i.e. pluripotency).

However, these approaches proved less successful in other inbred strain backgrounds. Indeed, some strains such as the nonobese diabetic (NOD) strain proved to be recalcitrant to ESC derivation (“nonpermissive”; Kawase et al., 1994; Gardner and Brook, 1997). This recalcitrance was ultimately surmounted by the adoption of “2i” culture conditions, which include small molecule inhibitors of ERK1/2 and GSK3 signaling (Nichols et al., 2009; Ying et al., 2008). It is now recognized that mESCs grown in 2i media exhibit a ground pluripotent state, characterized by homogeneous and stable expression of a core transcriptional pluripotency network resembling that of the pre-implantation epiblast (Abranches et al., 2014; Marks et al., 2012; Nichols et al., 2009; Ying et al., 2008). However, the stability of this state varies depending on genetic background. Hanna et al. (2009) reported that NOD mESCs exhibit metastability of the ground state in the absence of exogenous ERK1/2 and GSK3 inhibition. Despite these advances in our understanding of the genetic and molecular basis of pluripotency, we lack a detailed understanding of the extent of metastability across diverse strains and the underlying genetic differences that drive interstrain variation in derivation efficiency, as well as ground state pluripotency and differentiation capacity *in vitro*.

Here we leverage genetic diversity accumulated over ~500,000 years of evolution along the mouse lineage to study metastability of ground state pluripotency in mESCs. A large subset of *Mus musculus* diversity is captured by the eight parental strains of inbred mice used to develop the Collaborative Cross (CC) recombinant inbred mouse resource and complementary Diversity Outbred (DO) heterogeneous stock (Chesler et al., 2016; Churchill et al., 2004, 2012; Threadgill and Churchill, 2012). DO mice segregate >40 million genetic variants with relatively balanced founder allele frequencies and a simple population structure that enables high-resolution genetic mapping with relatively small sample sizes. To determine the genetic and molecular mechanisms leading to pluripotent ground state metastability, we derived mESCs from hundreds of DO mice and measured molecular phenotypes in cells grown in the absence of ERK2 inhibition.

## Results

### Genetic background drives variation in ground state pluripotency

To better understand the consequences of genetic background on the transcriptional networks governing the maintenance of pluripotency, we took advantage of germline-competent, euploid mESC lines that we previously derived from the eight genetically diverse inbred founder strains of the mouse CC (Czechanski et al., 2014). These include the classical laboratory strains C57BL/6J, A/J, 129S1/SvImJ, NZO/HILtJ, NOD/ShiLtJ (referred to here as B6, AJ, 129S1, NZO, and NOD, respectively); as well as inbred wild-derived strains representing three subspecies of *Mus musculus* including WSB/EiJ (WSB, *M.m. domesticus*), CAST/EiJ (CAST, *M.m. castaneus*), and PWD/PhJ (PWD, *M.m. musculus*; a consubspecific inbred wild-derived strain with similar geographic origin to the CC founder strain PWK/PhJ). We profiled genome-wide gene expression by RNA-Seq in three male mESCs from each strain. Principal component analysis of global transcriptional profiles revealed that inbred strain background is the major driver of transcriptional variability in these mESC lines (Figure 1A). Moreover, clustering by expression of core pluripotency genes and early lineage markers demonstrated heterogeneity across genetic backgrounds (Figure 1B). Although core pluripotency transcription factors like *Oct4/Pou5f1* and *Nanog* were highly expressed in all cell lines, expression of these and other genes varied by up to two orders of magnitude between cell lines (Figure 1B and 1C; Table S1), despite 2i culture conditions. Among the top differentially expressed genes (false discovery rate [FDR] = 5%; Table S1) were members of cytokine receptor signaling pathways, including leukemia inhibitory factor receptor (*Lifr*), where mESCs from wild-derived strains (CAST, PWD, and WSB) and the known recalcitrant classical laboratory strain NOD showed 1.5-fold to 5-fold reduced expression compared to B6, AJ, and 129S1.

**Figure 1.**
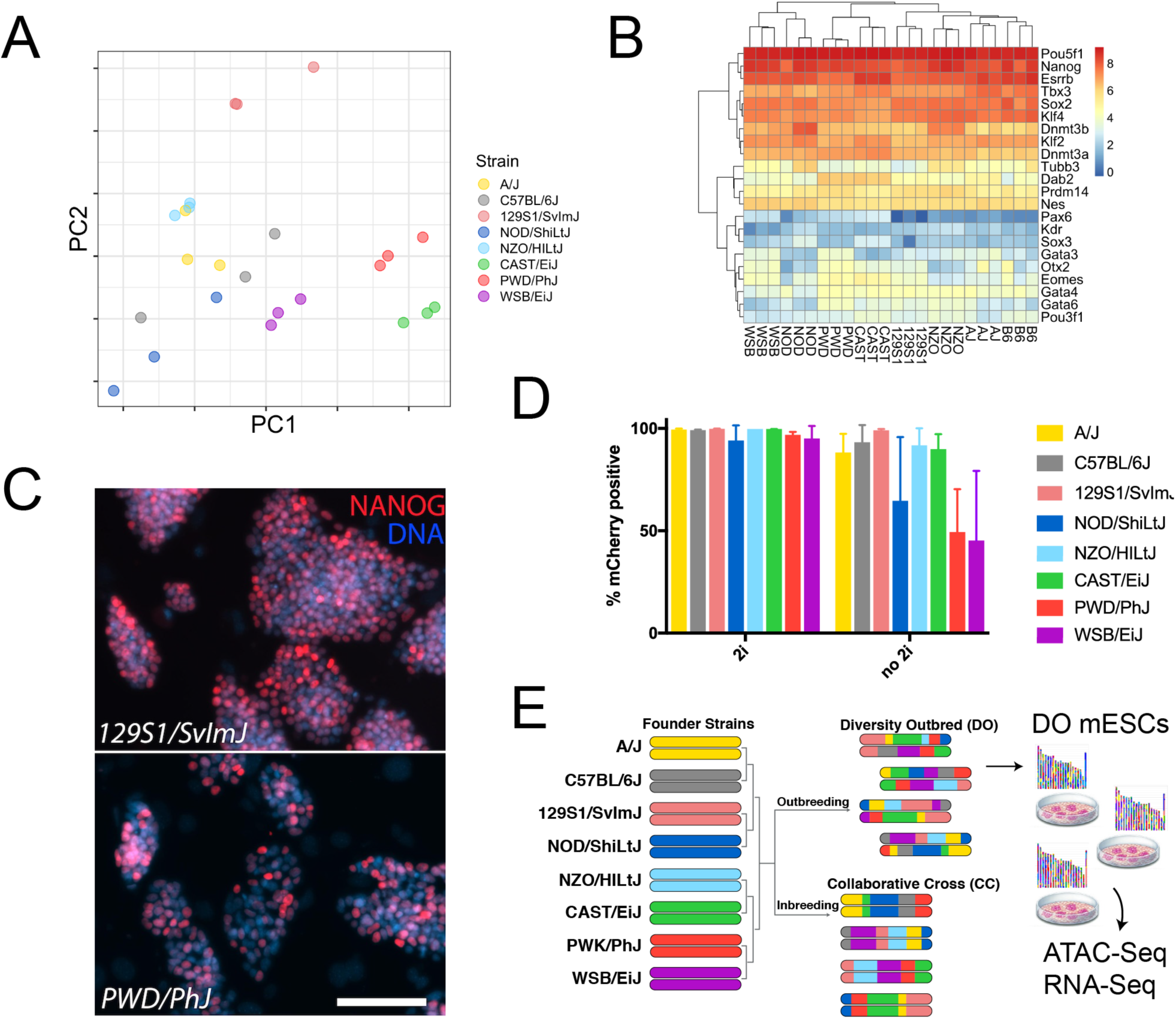
Molecular Indicators of the Pluripotent Ground State Vary in mESCs from Genetically Diverse Inbred Backgrounds. (A) Principal component analysis of global transcriptional profiles of mESCs from eight genetically diverse inbred backgrounds. Distinct dots from same genetic background represent biological replicate cell lines derived from the same inbred strain. (B) Expression of core pluripotency genes and early lineage markers shows both intrastrain (e.g. B6) and interstrain (e.g. NOD vs. WSB) variability. (C, D) Quantification of NANOG expression in mESC lines constructed from diverse inbred backgrounds using immunofluorescence and a *Nanog*-mCherry reporter knock-in. (C) Representative images of cells from two mESC lines that differ markedly in NANOG expression in 2i culture conditions. (D) Instability of *Nanog*-mCherry expression in knock-in cell lines in the absence of 2i in some inbred strain backgrounds, including known recalcitrant strains like NOD, as well as the wild derived strains PWD and WSB. (E) Overview of experimental design. Diversity Outbred mESCs segregate the bulk of variation that is fixed on inbred backgrounds in the strains shown in A-D and provide a powerful platform for genetic mapping of molecular QTL.

Based on this pattern of expression and previously published data (Silva et al., 2009), we predicted that the removal of ERK and GSK3 inhibition would destabilize *Nanog* expression, because these pathways act downstream of *Lifr* and are more proximal to the suite of genes that reinforce pluripotency. To test this, we introduced an in-frame, self-cleaving mCherry reporter fused to the stop codon of the endogenous *Nanog* locus (Yang et al., 2013) in each of the 24 inbred strain mESC lines. We found that in NOD, PWD, and WSB mESCs, fewer than two-thirds of cells expressed *Nanog* upon removal of ERK and GSK3 inhibition (Figure 1D), demonstrating that mESCs from these backgrounds depend on ERK and GSK3 inhibition to maintain stable *Nanog* expression even in the presence of exogenous LIF. This observation confirms earlier reports of ground state metastability in NOD (Hanna et al., 2009), and demonstrates that other genetically diverse strains show a similar ground state metastability phenotype.

### Diversity Outbred mESCs permit a genetic approach to dissecting variation in ground state pluripotency

To understand the genetic control of ground state pluripotency among mESCs, we leveraged the Diversity Outbred (DO) model (Figure 1E) by deriving ESCs from DO blastocysts. We selected 213 lines that met quality control criteria including absence of gross aneuploidies and availability of high-quality SNP genotypes. We expanded these mESCs under sensitized conditions in the absence of ERK inhibition to expose ground state metastability which we predicted would reveal genetic influences on this state (Ying et al., 2008). To characterize the molecular landscapes of our genetically diverse mESCs, we measured steady-state transcript abundance by RNA-Seq (*N* = 185 lines) and chromatin accessibility by ATAC-Seq (*N* = 192 lines) as an indicator of regulatory element activity. Both assays were performed in a common set of 166 cell lines.

We quantified expression of 15,185 genes and chromatin accessibility at 102,173 locations (peaks) across the DO mESCs. We found that 84% of transcripts (12,764) and 47% of chromatin accessibility peaks (47,786) showed variation that was significantly heritable (FDR = 5%). While all cell lines showed transcriptional hallmarks of pluripotency, including overall high expression of genes involved in regulation of pluripotency and low expression of lineage markers (Figure 2A), we observed variability in expression of these genes among cell lines. Variable activation of proximal gene regulatory elements (i.e. promoters, enhancers, insulators) may underlie variable gene expression and be detected by ATAC-Seq as changes in chromatin accessibility. As expected, we observe open chromatin at or near promoters (Figure S1A), and variable amplitude of those proximal peaks is highly correlated to gene expression differences across the DO mESC lines (orange/red diagonal line in Figure 2B).

**Figure 2:**
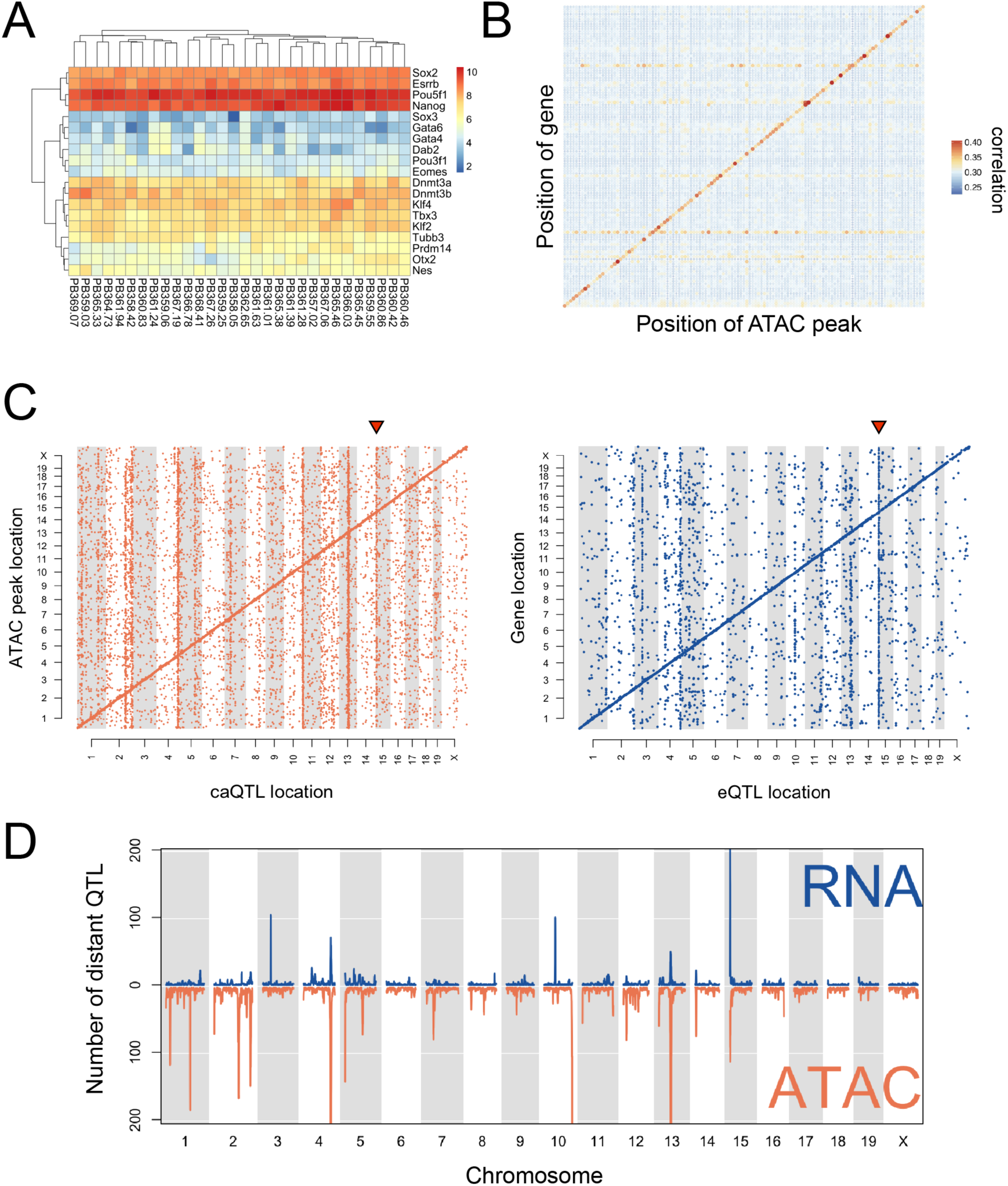
Genetic variation drives local and distant regulation of chromatin accessibility and gene expression. (A) DO mESCs show broadly high expression of core pluripotency genes and broadly low expression of early lineage markers. (B) Correlations between gene expression and chromatin accessibility across the genome. Only autosomal genes with a local eQTL (LOD >7.5) are shown. Features (genes and chromatin peaks) were grouped in 1cM bins. Correlations between all transcripts in the bin and accessibility of all chromatin peaks in the bin were computed. Points on plot are colored and sized according to the maximum correlation in the 1cM bin. Correlation tends to be highest between gene expression and local chromatin accessibility (diagonal). (C) caQTL and eQTL maps showing the genomic locations of significant QTL. Pronounced diagonal bands reflect the predominance of local caQTL and eQTL. Triangle above caQTL/eQTL hotspots on Chr 15 indicates the hotspot discussed in detail later in the main text. (D) Distinct and co-occuring caQTL and eQTL hotspots. Hotspots were defined as described in the main text. Distant QTL linkages are plotted along the same genomic coordinates for each data type (gene expression above; chromatin accessibility below) to facilitate comparison.

### Molecular mapping identifies extensive distal regulation of chromatin and gene expression

We conducted quantitative trait locus (QTL) mapping to understand the genetic regulation of chromatin accessibility and gene expression in mESCs. We mapped 33,196 chromatin accessibility QTL (caQTL) for 30,458 distinct chromatin accessibility peaks (Figure 2C, left; LOD > 7.5, corresponding to a permutation-based genome-wide *P* < 0.05) and 6,589 eQTL for 5,853 distinct genes (Fig 2C, right; LOD > 7.6, permutation-based genome-wide *P* < 0.05). Most QTL are local (69% of caQTL and 63% of eQTL) and evident as a clear diagonal band in Figure 2C; these local associations likely drive much of the observed high correlation between chromatin accessibility and transcript abundance of nearby genes (Figure 2B). Distant QTL are also numerous in both datasets. For example, we found that the transcription factor *Gata6,* a known marker of endoderm differentiation, had a local eQTL on Chr 18 largely driven by the PWK genotype. *Gata4* and *Sox17*, two additional markers of endoderm differentiation (Kanai-Azuma et al., 2002), have distal eQTL that map near *Gata6*, which we identified as the likely causal intermediate regulating transcript abundance of these genes (mediation analysis; see below). In agreement with this genetic variation in gene expression, mESCs derived from mice sharing the PWD *M.m. musculus* background more readily differentiate into certain endoderm lineages (Ortmann et al., this issue). We also identify multiple instances of loci that control chromatin accessibility and/or gene expression at hundreds of distal sites (Figure 2C). These clustered QTL, evident as vertical bands in Figure 2C (often referred to as QTL “hotspots”; Schadt et al., 2003), are of particular interest because they likely harbor regulatory genes (so-called “master regulators”) with widespread *trans*-acting effects on the regulation of many molecular features elsewhere in the genome. We identified thirteen such QTL hotspots (eight chromatin accessibility and five gene expression; on Chr 1, 2, 3, 4, 5, 10, 13, and 15), of which three were shared in both chromatin accessibility and gene expression data (Figure 2D). Although we observed aneuploidies in multiple DO mESC lines (Figure S2), hotspot signals persisted after controlling for possible aneuploidy (data not shown). Hotspot loci range in size from ~2-10Mb and contain an average of 106 annotated genes (minimum 34, maximum 249) representing candidate master regulators that may drive large-scale epigenetic or transcriptional changes (Table S2).

Next, we applied mediation analysis (see Methods; Chick et al., 2016) to identify the gene or genomic feature within each QTL hotspot that most likely conferred the observed QTL effect on the target genes or chromatin peaks. In many cases, the observed distant effect stems from local genetic variation within the QTL hotspot that directly influences the transcript abundance of a “mediator” gene (e.g. genetic variation → [mediator] → downstream targets). Mediation analysis of caQTL and eQTL hotspots narrowed our lists of putative master regulators between 4- and 59-fold. By choosing the best candidate mediator (largest LOD drop) of each target gene and imposing weak thresholds (candidate mediator must be best mediator for at least 5 hotspot targets and >5% of targets), we obtained 2-26 candidate mediators per hotspot (Table S3). Although mediation provides compelling statistical evidence for a candidate based on its expression, we cannot fully discount the possibility that another causal driver of the QTL hotspot acts through other mechanisms, such as an unannotated RNA, nonsynonymous coding variant, or inter-chromosomal chromatin contacts.

Mediation analysis of caQTL and eQTL highlighted several observations relevant to mESC biology. For example, an eQTL hotspot on Chr 3 (centered at ~51Mb) regulates the transcript abundance of 109 distant genes, many of which are involved in the chromatin remodeling SWI/SNF complex (16.8-fold enrichment; corrected *P* = 2.1×10^-4^). For 62 (57%) of these target genes, expression of the exosome complex subunit *Exosc8* was identified as the best mediator in this region. Exosome complex degrades AU-rich element-containing mRNAs and is associated with genome integrity and survival in human iPSCs (Skamagki et al., 2017). At a caQTL hotspot on chromosome 5 (centered at ~6Mb) regulating 141 distant peaks, the transcript coding for the multi-pass membrane protein *Steap2* was the best mediator for 75 (53%) distant targets. *Steap2* has been identified as a marker of mesenchymal stem cells and associated with the haematopoetic stem cell marker *Sca-1/Ly6a* (Vaghjiani et al., 2009), hinting that this gene may be involved in epigenetic changes leading toward lineage commitment.

We also found an eQTL hotspot on Chr 10 that influenced the expression of 100 genes. Interestingly, the list of target genes regulated by this locus demonstrates striking overlap with genes that are strongly and specifically upregulated in the rare 2-cell (2C)-like state (47/123; P < 1×10-16; Hendrickson et al., 2017). Cells in the 2C-like state are totipotent and present in mESC cultures derived from B6/129 substrain backgrounds at low frequency (~1%; Ishiuchi et al., 2015; Macfarlan et al., 2012). These cells are defined by their transcriptional similarity to totipotent early embryos, including characteristic activation of the endogenous retroviral element MERVL (Macfarlan et al., 2012). Expression of the MERVL long terminal repeat (MT2_mm) in DO mESCs is strongly correlated with expression of Chr 10 QTL target genes (□ = 0.96; Figure S1B). We found that abundance of the DUX transcript (*Duxf3*) was among the top two candidate mediators, although *Duxf3* locus is ~2Mb from the location of our mapped QTL. This is likely due to inconsistency in genome assembly at this chromosomal region, where there are known gaps in the reference assembly (GRCm38.p6) and reported segmental duplications (Lilue et al., 2018). These observations, in tandem with prior research demonstrating a role for DUX in activating 2C-like genes (Hendrickson et al., 2017), suggest that variation in *Duxf3* expression drives this hotspot. Rather than indicating differences in expression of 2C-like genes, this hotspot may reflect differences in cell state composition between DO mESC lines, with some lines having no/few cells in the 2C-like state and other lines having a relatively higher frequency of cells in this state.

### *Lifr* drives a suite of gene expression changes with functional consequences for mESCs

The QTL hotspot located on proximal Chr 15 (centered at ~8Mb; marked by inverted red triangles in Figure 2C) represents a putative master regulator that controls both caQTL and eQTL (Figure 2B). Furthermore, the suite of 254 target transcripts shows strong enrichment for genes known to have roles in maintaining pluripotency, including *Klf4, Lin28a*, and *Cxxc1*, highlighting the importance of this locus for understanding genetic differences in mESC biology. For the 254 eQTL target genes, estimates of founder haplotype effects at the Chr 15 locus indicated that cell lines carrying ancestry from classical laboratory strains (except NOD) tended to be similarly affected by the QTL genotype, while ancestry from NOD or the three wild-derived strains resulted in opposite effects (Figure 3B).

**Figure 3.**
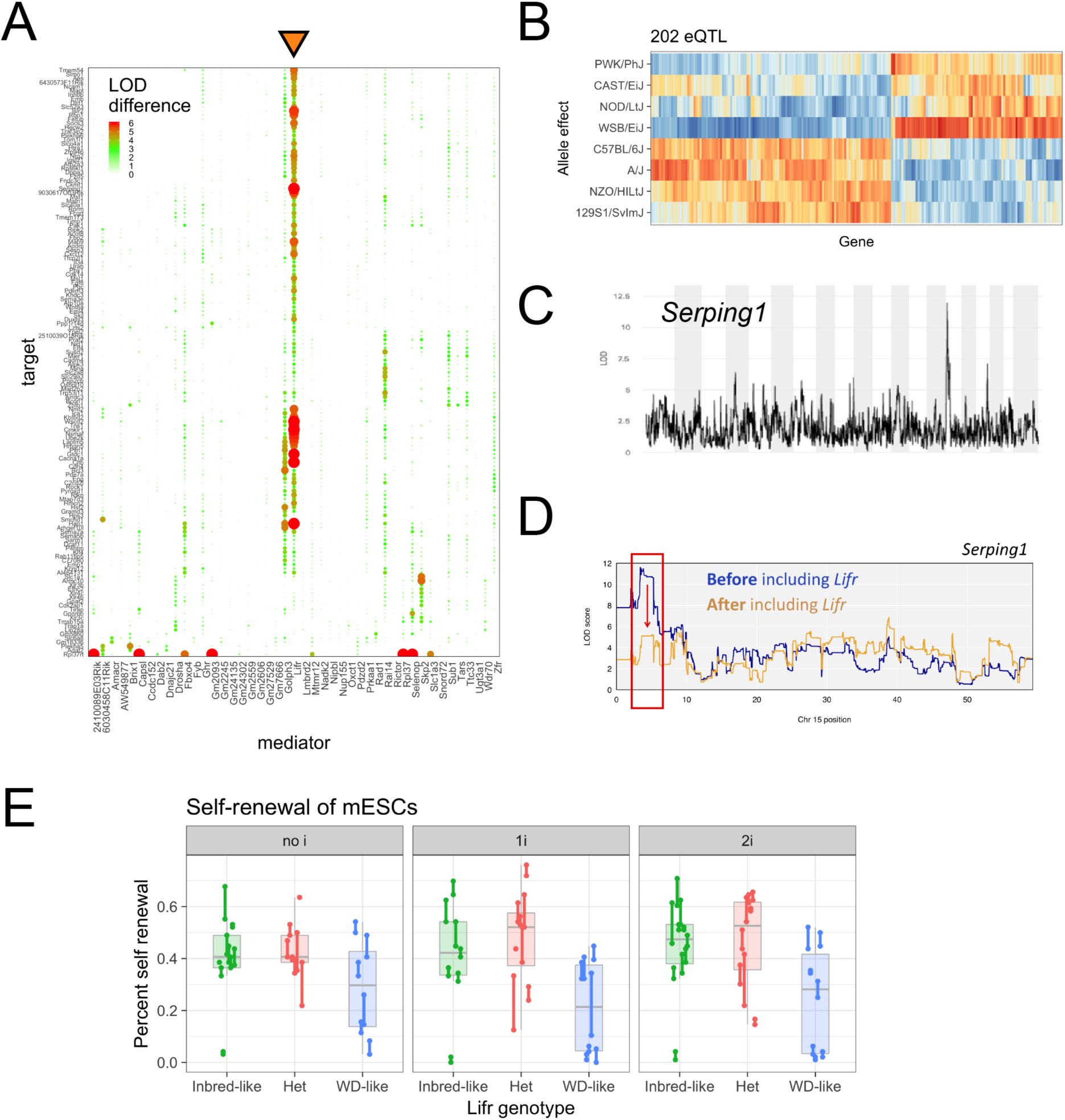
QTL mapping to chromosome 15 implicates the cytokine receptor *Lifr* as a key gene regulating chromatin and gene expression at genomically distant loci. (A) Mediation analysis reveals a single gene, *Lifr*, as the best mediator for 49% (122/254) of the distant gene expression targets (*Lifr* result is shown in the column marked with an inverted orange triangle). Dotplot shows candidate mediators located within Chr 15 hotspot along *x*-axis, and targets of the hotspot on *y*-axis. The LOD difference for each target gene after including each candidate mediator is plotted as a dot with size and color proportional to the drop in LOD (higher drop indicating better mediation; bright red indicates drop of ≥6; white indicates drop of ≤0). (B) Allelic effects for targets of the Chr 15 hotspot. Higher effects (red/orange) indicate higher expression of the target gene in mESCs carrying a haplotype derived from the founder listed at left. Lower effects (blue/light blue) indicate the opposite. For each target gene, effects are scaled to have mean 0 and variance 1. (C, D) *Serping1* provides a representative example of a gene whose expression is consistent with mediation by *Lifr* expression. (C) *Serping1* is located on chromosome 2 and has a strong distal eQTL (LOD = 12) on Chr 15. (D) When *Lifr* gene expression is included as a covariate in the genetic mapping model for *Serping1* expression, the LOD significance score for the Chr 15 QTL drops precipitously (>6 LOD units). (E) mESC self-renewal percentage stratified by *Lifr* genotype. Measurements were undertaken in three media conditions shown in facets and labeled at top. Replicate measurements within each panel are connected by a line.

Mediation analysis revealed a single gene, *Lifr*, as the best mediator for 48% (122/254) of the distant gene expression targets (Figure 3A; results of testing *Lifr* as a mediator are presented in the column marked with an inverted orange triangle). *Serping1* provides a representative example of a gene whose expression is consistent with mediation by *Lifr* expression. *Serping1* is located on Chr 2 and has a strong distal eQTL (LOD = 12) on Chr 15 (Figure 3C). When *Lifr* gene expression is included as a covariate in the genetic mapping model for *Serping1* expression, the LOD significance score for the Chr 15 QTL drops precipitously (>6 LOD units; Figure 3D), suggesting that changes in *Lifr* gene expression fully capture and confer the observed distant eQTL effect on *Serping1 (i.e. Chr 15 QTL* → *[Lifr]* → *[Serping1])*. QTL mapping of *Lifr* transcript abundance revealed a strong local eQTL, suggesting a *cis-*regulatory variant that directly influences *Lifr* expression and leads to shifts in the expression of downstream genes.

To validate the effect of *Lifr* gene expression on its downstream genes in an orthogonal genetic system, we derived mESCs from F1 intercrosses of recombinant inbred CC lines (CC-RIX) segregating the same genetic variation as DO mice. We quantified the expression of five Chr 15 eQTL target genes that were predicted to be mediated by *Lifr* expression (*Rbp1, Cxcl12, Hap1, Klf4*, and *Socs3*) by qRT-PCR in CC-RIX mESCs. The expression of each of these genes was correlated with *Lifr* expression in the CC-RIX lines, and both the magnitude and direction of this correlation agreed with our results in the DO mESCs (Figure S3), supporting our prediction that *Lifr* is the primary causal mediator of genes in the Chr 15 QTL hotspot.

Self-renewal is an important property of pluripotent cells that is lost upon lineage commitment. To examine the functional significance of *Lifr* genotype in the context of mESC pluripotency, we tested the ability of CC-RIX lines to proliferate from single cells in colony forming assays as a quantitative measure of self-renewal. Categorizing the hybrid CC-RIX lines on the basis of their parent CC genotypes at the *Lifr* Chr 15 locus (Table S4), we found that *Lifr* genotype was a good predictor of self-renewal with lines homozygous for the wild-derived-like *Lifr* allele showing abrogated self-renewal (Figure 3E). Thus, genotype at the *Lifr* locus influences a suite of downstream gene expression changes that collectively modulate cell pluripotency.

### A single nucleotide variant influences ground state pluripotency through *Lifr*

In order to identify causal genetic variants within the Chr 15 QTL hotspot, we searched for single nucleotide variants (SNVs), insertions, or deletions that matched the 4:4 strain pattern seen in targets of this QTL (Figure 3B). Although there are over 20,000 segregating genetic variants within the 1.5Mb region surrounding *Lifr*, only 185 matched this unusual pattern, where NOD shared the same allele as the three wild-derived strains (CAST, PWK, and WSB). Given that *Lifr* shows a local eQTL, and our observation that local chromatin accessibility correlates with local gene expression (Figure 2B), we used variation in regions of open chromatin within +/− 2Mb surrounding *Lifr* to identify putative regulatory elements. The best-mediating chromatin accessibility peak was located ~10kb upstream of the transcriptional start site of *Lifr,* and overlaps a DNaseI hypersensitive site that is unique to mESCs (Figure 4A; blue tracks show mESCs). Critically, a single SNP (rs50454566, chr15:7116944 [GRCm38/mm10], T/A) with the unusual 4:4 pattern lies at the apex of this chromatin accessibility peak, and DO strains homozygous for the wild-derived-like allele (A/A) have lower chromatin accessibility and *Lifr* expression compared to strains homozygous for the laboratory strain-like allele (T/T) (Figure 4B). Founder inbred strain transcriptome data (2i) and qRT-PCR from our 1i (LIF + GSK3 inhibition) conditions confirmed the correlation between strain genotypes at this SNP and *Lifr* expression, where the classical laboratory strain genotype (T/T) and the wild-derived-like genotype (A/A) confer high and low expression, respectively (Figure S4A and S4B).

**Figure 4.**
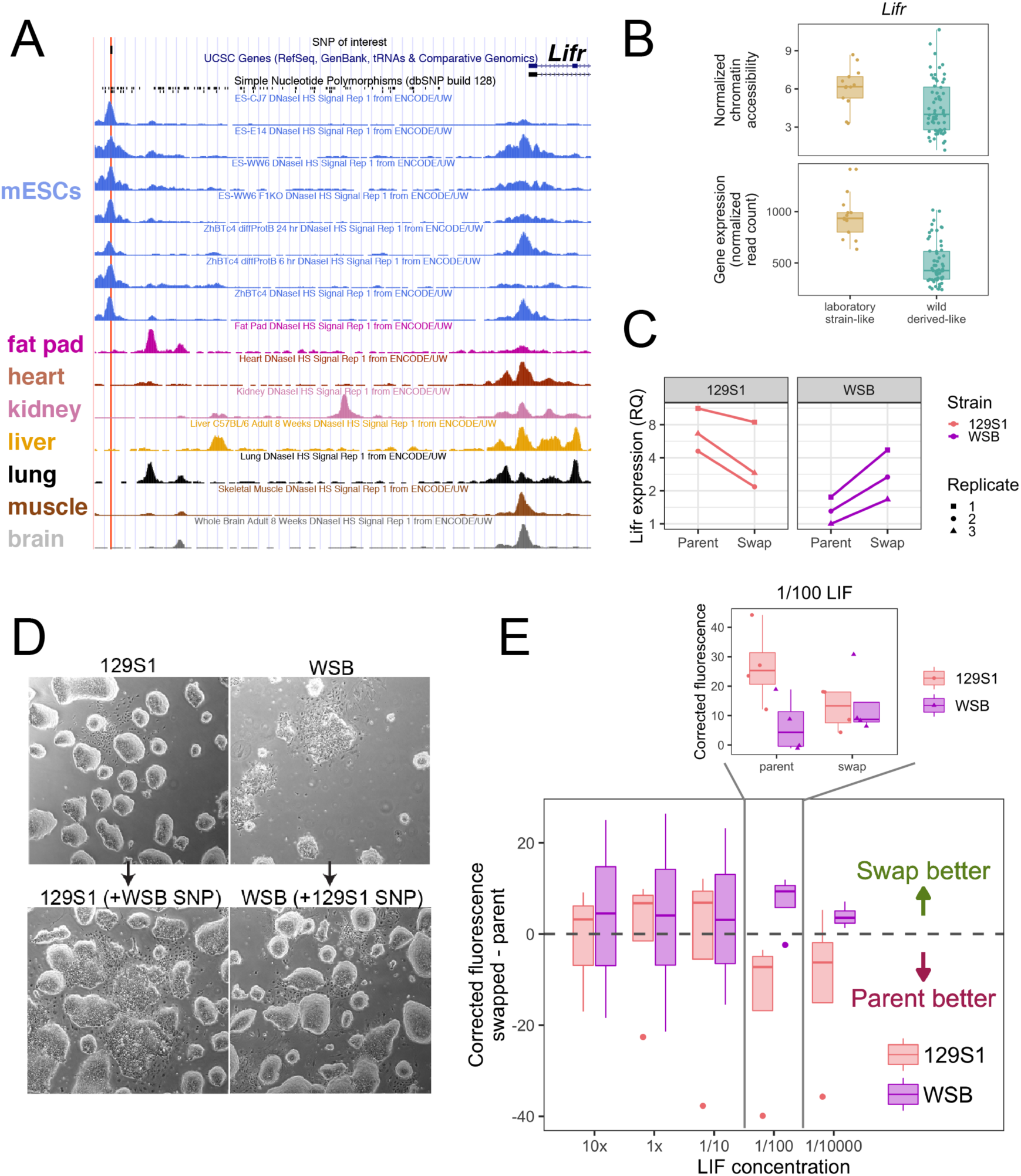
A single variant upstream of *Lifr* influences mESC pluripotency. (A) 15kb surrounding the *Lifr* transcriptional start site. Top track shows the SNP of interest discussed in the main text; dbSNP: rs50454566, chr15:7116944 (GRCm38/mm10), T/A. Lower tracks show DNaseI hypersensitive site data from the mouse ENCODE project (Mouse ENCODE Consortium et al., 2012). (B) DO strains homozygous for the wild-derived-like allele (A/A) have lower chromatin accessibility and *Lifr* expression compared to strains homozygous for the laboratory strain-like allele (T/T). (C) Under 2i growth conditions, *Lifr* expression is higher in WSB mESC clones that harbor the classic laboratory inbred strain allele (T/T; “swap”) with respect to the parental WSB mESC, and lower in 129S1 mESC clones that harbor the wild-derived-like allele (A/A; swap) with respect to the parental 129S1 line. Quantitative RT-PCR data for three replicate experiments are shown. (D) Colony morphology of WSB mESC clones harboring the classic inbred strain allele improves under 2i growth conditions. The opposing effect was somewhat less consistent in 129S1 mESC clones harboring the wild-derived-like allele. (E) mESC lines grown in the absence of ERK and GSK3 beta inhibition depend on cytokine signaling and exogenous LIF to maintain robust *Nanog* expression. The classic inbred laboratory strain SNP improved the WSB mESC responsiveness to limiting concentrations of LIF (1/100) over the parental WSB ESC lines, whereas the wild-derived-like SNP degraded the responsiveness of the 129S1 mESC lines (N=4 experiments).

To validate this SNP as a causal variant underlying the observed effects on *Lifr* expression and ground state pluripotency, we selected two founder strain *Nanog*-mCherry mESC lines with opposing SNP genotypes (129S1 [T/T] and WSB [A/A]) and performed reciprocal allele swaps to measure the effect of this SNP on *Lifr* expression and pluripotency. Placing the wild-derived-like variant on an otherwise 129S1 genetic background (129S1^[A/A]^) reduced *Lifr* transcript abundance (Figure 4C) and *Nanog-*mCherry expression (Figure S4C), whereas placing the common laboratory variant on an otherwise WSB genetic background (WSB^[T/T]^) increased *Lifr* and *Nanog*-mCherry expression (Figures 4C and S4C). This allele swap had a striking impact on colony morphology, with replacement of the single SNP upstream of *Lifr* correcting the characteristically poor colony morphology of WSB mESCs (Figure 4D). We further quantified the percentage of pluripotent cells (cells expressing *Nanog-* mCherry) across a range of LIF concentrations in the absence of 2i to test the robustness of cytokine signaling in these cell lines. Compared to 129S1 mESCs, 129S1^[A/A]^ mESCs show reduced sensitivity to exogenous LIF and lower levels of pluripotency as evidenced by lower *Nanog*-mCherry in limiting concentrations of LIF (1/100 and 1/10000; Figure 4E). In contrast, WSB^[T/T]^ mESCs exhibit higher levels of pluripotency relative to WSB mESCs, with 129S1-like expression of *Nanog-*mCherry at the same limiting concentrations of LIF (Figure 4E). Thus, a single nucleotide influences chromatin accessibility upstream of *Lifr*, which alters *Lifr* expression and leads to downstream changes in a suite of genes with effects on pluripotency.

## Discussion

Genetic background strongly influences molecular phenotypes in ESCs, and can be harnessed to discover drivers of differences in ESC biology. These molecular phenotypes are reflections of cell states that occur along a developmental continuum. Our genetic approach identifies multiple loci influencing metastability of ground state pluripotency, and provides mechanistic details clarifying the recalcitrance of some inbred mouse strains to ESC derivation in the absence of ERK1/2 and GSK3 inhibition. In particular, natural variation plays a critical role in shaping the ability of mESCs to respond to exogenous signals, explaining why bypass of cytokine signaling through ERK1/2 and GSK3 inhibition stabilizes ground state pluripotency in mESCs from those strain backgrounds that carry the wild-derived-like *Lifr* allele. It is likely that genetic differences in responses to exogenous cues drive other aspects of mESC biology; for example, differences in *in vitro* differentiation capacity (Ortmann et al., this issue) may partially reflect variable responses to differentiation cues originally optimized for a single genetic background.

The Chr 15 QTL hotspot and causal *Lifr* upstream regulatory SNP only partially explain the transcriptional and epigenetic variation present in DO mESCs cultured in the absence of ERK1/2 inhibition. Our data reveal additional hotspot loci that likely impact ground state metastability and other cell states (e.g. Chrs 3, 5, and 10). For example, the Chr 10 QTL hotspot regulates many genes expressed in the early totipotent/2C-like cell state (Macfarlan et al., 2012), suggesting variation in the abundance of this rare and transient state among our DO mESCs, which has intriguing implications for the capture of cells that more persistently display their expanded developmental potential (Baker and Pera, 2018). Further detailed studies of the causal variants underlying these QTL hotspots, as well as studies incorporating improved genome assemblies (Lilue et al., 2018) and high-resolution single cell profiling to disentangle heterogeneity of cell states, will reveal genes and pathways driving intraspecies variation in pluripotency and differentiation capacity.

Genetic background also strongly influences molecular phenotypes and differentiation capacity in human ESCs and induced pluripotent stem cells (hiPSCs). Differences between genetic backgrounds are a dominant source of inter-line variability in hESCs and hiPSCs (Burrows et al., 2016; Carcamo-Orive et al., 2017; Choi et al., 2015; DeBoever et al., 2017; Féraud et al., 2016; Kajiwara et al., 2012; Kyttälä et al., 2016; Osafune et al., 2008; Ramos-Mejia et al., 2010). QTL mapping in panels of differentiated hiPSCs has revealed loci controlling transcript abundance and chromatin accessibility in lineage-committed cells (Alasoo et al., 2018; Schwartzentruber et al., 2018). There are significant differences between mESCs and hESCs; in light of our results it is notable that LIF is dispensable for the maintenance of hESCs (Thomson, 1998). Nevertheless, compared to mESCs, hESCs are functionally and phenotypically more similar to mouse EpiSCs isolated from post-implantation blastocyts (Brons et al., 2007; Tesar et al., 2007), which also do not require LIF. Given reports that hESCs can be coerced into a more naïve pluripotent state by ectopic expression of pluripotency factors including LIF (Buecker et al., 2010; Hanna et al., 2010), as well as the role of LIF signaling in blastocyst implantation in humans (Aghajanova, 2004), it is likely that shared regulatory circuits involving LIF and *Lifr* contribute to transcriptional changes occurring at these early developmental timepoints in both species. The unique ability of diverse mouse ESCs to provide access to very early developmental states coupled with unbiased and high-resolution mapping of gene regulatory relationships underscores the utility of carefully designed model organism genetic resources for basic biological discovery.

## Acknowledgments

We thank David Aylor and Thomas Konneker for their early contributions in the analysis of a subset of the inbred strain transcriptome data. We thank Steve Murray, Kevin Peterson, and Susan Kales for graciously sharing the mCherry reporter construct. We thank Martin Pera and Daniel Cortes-Perez for constructive feedback on this project and the manuscript. We thank Richard Paules and Kristine Witt for scientific contributions toward development of the DO mESCs and Alex Merrick of NIEHS for reviewing the initial manuscript drafts. Funding sources included HHSN273201500196P (NIEHS-NTP to TC), OD010921 (LGR), OD011102 (LGR), GM070683 (GAC) and The Jackson Laboratory Director’s Innovation Fund (CLB, SCM, LGR, GAC, DAS, AC, IG, NR). The University of Virginia Genome Analysis and Technology Core provided sequencing services for DO RNA-Seq. The Jackson Laboratory Genome Technologies Service (JAX-GT) provided sequencing services for founder mESC RNA-Seq and DO ATAC-Seq data generation. JAX-GT are supported by the National Cancer Institute under award number P30CA034196.

## Author Contributions

Conceptualization, LGR, SCM, CLB, DAS and GAC; Methodology, DAS, LGR, SCM, CLB, TC, AC, Resources, SD and WQ; Investigation, AC, CByers, CS, DKP, MP, CO, WM, IG, LGR, AC, WM, IG; Data curation and formal analysis, DAS, SM, CLB, SA, NR; Visualization, DS; Writing - original draft, review, and editing, DS, LR, SM, SA, CLB, TC, GC; Supervision, LGR, CLB, SM; Project administration, LGR, CLB, SCM, GAC, Funding acquisition, TC, LGR, CLB, SCM, DAS, GAC.

## Methods

### Founder inbred strain mESCs

#### Derivation and culture

Euploid (>70%), germline competent, male mESCs were derived from C57BL/6J (JR# 000664), 129S1/SvImJ (JR#002448), A/J (JR#000646), BALB/cByJ (JR#001026), NOD/ShiLtJ (JR#001976), NZO/HlLtJ (JR#002105), CAST/EiJ (JR#000928), PWD/PhJ (JR#004660), WSB/EiJ (JR#001145) as previously described. Note that PWD/PhJ is distinct from PWK/PhJ, which is a Collaborative Cross founder strain. Nevertheless, both are inbred strains of the *M. m. musculus* subspecies that are derived from wild mice trapped near Prague, Czech Republic. Derivation and characterization (mycoplasma testing, SNP genotyping, pluripotency marker expression, chromosome counting, germline testing) of the mESCs were as previously described (Czechanski et al., 2014). For each strain we initially derived 15-40 unique mESC lines from individual blastocysts and adapted lines to ESM, 2i+LIF (“2i”) media in the presence of mitotically inactivated mouse embryonic fibroblasts (MEFs, C57BL/6J) (ESM, 2i+LIF: Dulbecco’s Modified Eagle Medium (DMEM) supplemented with 15% fetal bovine serum, 100 U/mL Penicillin-Streptomycin, 2mM GlutaMAX, 0.1mM non-essential amino acids, 1mM sodium pyruvate, 0.1mM 2-mercaptoethanol, 500pM LIF, 1uM PD0325901, and 3uM CHIR99021). Of these, three male lines from each genotype were selected for germline testing on the basis of robust expression of pluripotency markers and euploidy. We validated the ability of these ESCs to contribute to the germline *in vivo*, the gold standard for functionally defining pluripotency in mESCs. The resulting 24 germline-competent, euploid mESC lines (Supplementary Table 1) provide a panel of genetically distinct mESCs that differ at >40 million sites across the genome.

#### RNA-Seq

Prior to harvesting cells (P6-P8) for RNA collection, MEFs were removed through sequential plating (2X, 1 hour) onto gelatin-coated dishes. RNA was harvested using the RNeasy (Qiagen) RNA extraction kit. Poly(A) RNA-seq libraries were constructed using the TruSeq Stranded mRNA Library Prep Kit (Illumina), including the addition of unique barcode sequences. Libraries were pooled and sequenced 125 bp paired-end on the HiSeq 2000 or 2500 (Illumina) using TruSeq SBS Kit v4 reagents (Illumina). To reduce spurious alignments, we constructed strain-specific genomes for the 8 inbred strains using the software tool g2gtools and genetic variation data from Mouse Genomes Project SNP and indel release version 5 (ftp://ftp-mouse.sanger.ac.uk/REL-1505-SNPs_Indels/). We used Ensembl gene annotation version release-84 to extract strain-specific transcriptomes and annotations for these strains (ftp://ftp.ensembl.org/pub/release-84/gtf/mus_musculus). RNA-Seq reads from each strain were aligned to the strain-specific transcriptomes using bowtie (Langmead et al., 2009) aligner with parameters “--best --strata -a -m 100 -v 3”. These settings retain all read alignments with the best alignment score allowing up to 3 mismatches for further analysis. We used EMASE (Raghupathy et al., 2018) to quantify isoform-level and gene-level expression abundances. EMASE uses an EM algorithm to accounting for reads with multiple alignments and estimated read counts for isoforms and genes. Read counts were estimated in this manner on both raw and batch corrected data. Further analyses were conducted on both raw and batch corrected read count data to identify and mitigate batch effects. We used DESeq2 (Love et al. 2014) to test for variation in gene expression driven by genetic background.

#### Nanog-mCherry engineering

A CRISPR/cas9 donor construct containing a Nanog-2A-mCherry reporter cassette designed and constructed by Yang et al. (2013) was obtained from Addgene (Plasmid #48680). Targeting arms were shortened to 1kb so that the updated vector extended from Chr6:122,712,552-122,714,555 (GRCm38/mm10) with the insert at Chr6:122,713,552 (GRCm38/mm10). A guide targeting the last coding exon of *Nanog* (CCACTTTATACTCTGAATGC) was cloned into the pSpCas9(BB)-2A-Puro (PX459) V2.0 (Addgene #62988). 5×10^5^ cells seeded on a 12-well plate were co-transfected with equimolar amounts (0.5ug each) of donor and guide vector using Lipofectamine 3000 (Invitrogen #L3000) reagent. Following puromycin selection, positive knock-in cells were isolated by mCherry reporter expression using a FACSAria flow sorter and single mCherry expressing cells were plated individually in 96 well plates for clonal expansion. Standard and long range PCR were used to confirm targeted knock-in (KI) of the reporter cassette. Homozygous *Nanog-mCherry* KI cell lines or heterozygous KI cell lines with intact WT *Nanog* alleles were selected using these molecular data. Functional NANOG protein expression was confirmed by flow cytometry.

#### Nanog-mCherry expression analysis

For each of the 8 Collaborative Cross founder strains, three independent *Nanog*-mCherry clones were passaged from ESM+2i (“2i”) conditions onto 24 well MEF plates in ESM+2i (“2i”) and in ESM (“no i”) for 48 hours with daily media changes until sub-confluent. An additional well with a non-targeted mESC line was plated in both conditions to serve as a negative control for flow cytometry. Wells were then harvested, resuspended in 2 ml of PBS with DAPI and processed on the FACSymphony FlowJo version 10 was used for analysis and the background fluorescence of the negative control was used to adjust for background fluorescence. Immunolabeling and flow cytometry were used to correlate *Nanog-mCherry* expression with NANOG protein expression as previously described (Reinholdt et al., 2012).

### Diversity Outbred mESCs

#### Derivation

Male and female Diversity Outbred mice (JR #009376) were obtained at approximately four weeks of age and maintained at Predictive Biology, Inc. for several rounds of breeding. At 24-26 days of age females were superovulated and subsequently mated to 7-15 week old males. mESC lines were derived from random blastocysts using previously described protocols (Czechanski et al., 2014).

#### Cell culture

Blastocysts were transferred to 2i medium in 96 well round-bottom ultra-low attachment plates for 5-7 days. Those that had some inner cell mass outgrowth were dispersed and transferred onto MEF feeders in 96 well flat-bottom tissue culture plates in ES medium (ESM, 1i+LIF: Dulbecco’s Modified Eagle Medium (DMEM) supplemented with 15% fetal bovine serum, 100 U/mL Penicillin-Streptomycin, 2mM GlutaMAX, 0.1mM non-essential amino acids, 1mM sodium pyruvate, 0.1mM 2-mercaptoethanol, approximately 2000U/ml LIF, and 3uM CHIR99021). Recombinant LIF protein was produced using a Chinese Hamster Ovary cell line. As the ES cells expanded, they were transferred into 24 well plates, followed by 6 well plates, and finally 10cM dishes, all without feeder cells but in the same media. Thus, as the cells expanded they were weaned off feeder cells by dilution.

#### Genotyping of mESCs

Cell lines were genotyped by Neogen Corp. (Lincoln, NE) using the Giga Mouse Universal Genotyping Array (GigaMUGA; Morgan et al., 2016). We used functions available in the argyle R package (Morgan et al., 2015) for quality control purposes. Specifically, we examined plots of B-allele frequency (BAF) and log2 intensity ratio (LRR) to scan for gross aneuploidies and did not move forward with any lines presenting such anomalies. We used the hidden Markov model implemented in DOQTL (Gatti et al., 2014) for haplotype reconstruction to obtain diplotype probabilities suitable for quantitative trait locus mapping.

#### RNA-Seq data generation - DO mESCs

Total RNA was isolated from each of 183 DO mESC lines and quantitated by paired-end RNA sequencing. Briefly, for each mESC line, one 15cm dish of cells was grown to near confluence, washed 3x with PBS, and mechanically harvested to produce 80-150mg wet weight cell pellets (~50M cells). About 100k cells from each frozen cell pellet were used for RNA sequencing. Next, total RNA was extracted using the Quick-RNA 96 well format kit (Zymo Research) with in-column DNase treatment. Sequencing libraries were prepared by Akesogen using the TruSeq Stranded mRNA HT kit (Illumina, Cat no. 20020595) and included ribosomal RNA reduction and poly-A selection, enzymatic fragmentation, cDNA synthesis from random hexamer priming, adapter ligation and PCR amplification steps to generate indexed, stranded mRNA-seq libraries. Libraries were checked for quality and quantitated with the Agilent Bioanalyzer, and samples that failed QC were repeated starting from the cryovial stage. Finally, pooled libraries were sequenced on the NextSeq platform (Illumina) using the NextSeq 500/550 High Output v2 150-cycle kits (Illumina, Cat no. FC-404-2002). To minimize technical variation, samples were randomly assigned to lanes prior to sample processing steps, barcoded, and multiplexed at 16 samples per flow cell, yielding 6M-55M 2×75bp paired-end (PE) reads per sample. Two of the 183 lines were grown in replicate, resulting in a total of 185 samples with RNA-Seq data. For downstream genotyping, quantitation, and eQTL mapping analyses (see below), only the first read of the pair was used.

#### ATAC-Seq data generation

To measure chromatin accessibility, ~100,000 cryopreserved DO mESCs were used in the Fast-ATAC protocol (Corces et al., 2016). Cryopreserved cells were thawed in a 37°C water bath, an aliquot of ~100,000 cells removed, then spun and washed with cold 50 *µ*l PBS to remove DMSO. The cell pellet was resuspended in the 50 *µ*l transposase reaction mix (25 *µ*l of 2X TD buffer, 2.5 *µ*l of TDE1, 0.5 *µ*l of 1% digitonin, and 22 *µ*l of nuclease-free water). Transposition was carried out for 30 min at 37°C, and DNA purified using a Qiagen MinElute kit. Libraries were amplified for a total of 9 cycles and purified using 1.7X AMPure beads. Nucleosome banding was visualized using the Agilent Tapestation. Libraries were subject to 100 bp single-end sequencing on an Illumina HiSeq 2500.

#### RNA-Seq data analysis

We aligned single-end 75bp reads with bwa v1.0.0 (Li and Durbin, 2009) to a pooled “8-way” transcriptome containing strain-specific isoform sequences from all eight DO founder strains. To construct the 8-way founder transcriptome, strain SNVs and short indels (release 1505) were downloaded from the Sanger Mouse Genomes Project (Keane et al., 2011) and incorporated into annotated transcripts (Ensembl release 82) using g2gtools v0.1.31. We inferred sample genotypes genome-wide from the RNA-Seq data using gbrs v0.1.6 (http://churchill-lab.github.io/gbrs/), and compared gbrs-derived genotypes to our DNA (GigaMUGA) genotypes to identify potential sample mix-ups. Typically, correlations between haplotype probabilities inferred from gbrs versus GigaMUGA on the same sample were on the order of 0.8-0.9, while correlations for these probabilities measured on different samples were <0.5. Using this procedure, we identified 10 DO mESC lines with incongruent genotypes, and were able to resolve nine of these errors. For the sample where a definitive genotype could not be ascertained, the gbrs genotype was used for QTL mapping.

We applied EMASE v0.10.16 (Raghupathy et al., 2018) to resolve multi-mapping reads and estimate transcript- and gene-level abundance for each sample. We filtered out genes for which the median TPM (transcripts per million) value was <0.5 or where more than half of the samples were zero (i.e. not expressed). Next, we normalized gene-level counts to the upper quartile value in each sample to account for differences in library size, and then used ComBat (Leek et al., 2012) to ameliorate any potential batch effects stemming from library preparation. Finally, we transformed upper quartile normalized, ComBat-adjusted values to rank normal scores using the ‘rankZ’ function in the DOQTL R package (Gatti et al., 2014).

#### eQTL mapping

We performed eQTL mapping on normalized, transformed gene-level expression values described above using the ‘scan1’ function in r/qtl2 (Broman et al., 2018). We included sex as an additive covariate in our mapping model. To assess genome-wide significance, we applied a permutation strategy (1,000 permutations). Using this approach, we established a cutoff of LOD >7.5 for reporting significant eQTL.

#### ATAC-Seq data analysis

We used Trimmomatic version 0.33 (Bolger et al., 2014) to trim Illumina adapters from 100bp ATAC-Seq reads with the following settings “ILLUMINACLIP:NexteraPE-PE.fa:2:30:7 TRAILING:3 SLIDINGWINDOW:4:15 MINLEN:36”. We mapped trimmed reads to the mouse reference genome (GRCm38) using bwa version 0.7.9a (Li and Durbin, 2009). For each sample, we filtered out secondary alignments and reads with mapping quality equal to zero, and ran MACS version 1.4.2 (Zhang et al., 2008) on the remaining alignments with the following settings “-f BAM -g mm -p 1e-5”. We used the rtracklayer version 1.34.2 R package (Lawrence et al., 2009) to read in peaks called by MACS across each chromosome, saving only those genomic windows that were contained within peaks in at least 5 samples in order to define peaks that are unlikely to be spurious (yet allowing for the possibility of peaks that are unique to only some samples). To reduce the incidence of clusters of narrow peaks, we merged peaks <20bp in width into the nearest peak within 100bp. This set of peaks constituted our consensus set; we then used bedtools version 2.26.0 (Quinlan and Hall, 2010) to quantify ATAC-Seq read depth across the peaks in each sample. We removed peaks that did not exceed one count per million in at least 20 samples (<1% of peaks), and normalized at the sample level using the TMM method implemented in the edgeR version 3.16.5 R package (Robinson et al., 2010). Finally, we transformed normalized values to rank normal scores using the ‘rankZ’ function in the DOQTL R package (Gatti et al., 2014).

#### caQTL mapping

We performed caQTL mapping on normalized, transformed gene-level expression values described above using the ‘scan1’ function in r/qtl2 (Broman et al., 2018). We included sex and sequencing plate as additive covariates in our mapping model. To assess genome-wide significance and correct for multiple testing, we applied a permutation strategy (1,000 permutations). Using this approach, we established a cutoff of LOD >7.6 for reporting significant caQTL.

#### Defining caQTL and eQTL hotspots

We defined distant QTL as QTL where the location of the peak was greater than 10Mb from the genomic feature being mapped. To define hotspots, we started with lists of distant caQTL and eQTL with LOD scores exceeding a genome-wide permutation-based threshold (*P* < 0.05; LOD 7.5 for eQTL and 7.6 for caQTL). We tallied up distant links within overlapping 1cM bins (0.25cM shift) across the genome. We selected the top 0.5% of bins with most distant links (for each data type) and defined these loci as hotspots. We collapsed adjacent bins into a single region to obtain coordinates of hotspot loci. We used PANTHER (Mi et al. 2016) to perform gene ontology enrichment tests of hotspot targets.

#### Mediation analysis

We used the ‘intermediate’ package in R (https://github.com/simecek/intermediate) to perform mediation analysis to identify transcripts and regions of open chromatin in that region that were likely to be the causal mediator of distant eQTL and caQTL. Briefly, for a given distant QTL, we first identified all expressed genes and chromatin peaks within 10Mb of the peak SNP. We then included the transcript abundance (or peak intensity) of these candidate mediators individually as additive covariates in the QTL mapping model, and compared LOD scores at the peak distant SNP with and without the addition of this covariate. In cases where the distant QTL effect is mediated by the chromatin state or transcript abundance of a gene near the locus, inclusion of that parameter as an additive covariate in the mapping model abolishes or significantly decreases the distant QTL effect, as evidenced by a decrease in LOD score. We calculated LOD scores using the ‘double-lod-diff’ method in r/intermediate to minimize the effects of missing data in our RNA-seq and ATAC-seq data sets.

#### Quantitative RT-PCR

RNA was extracted from 1×10^6^ mESCs according to the manufacturer’s instructions using RNeasy Plus Mini Kit (Qiagen, cat. No. 74134) after cellular homogenization using QIAshredder (Qiagen, cat. No. 79654). RNA was eluted in 50*µ*l RNase free water and quantified with Nanodrop. cDNA was generated using High Capacity RNA to cDNA Kit (Thermofisher, cat. No. 4387406) with 500ng of RNA per sample. Resulting cDNA was diluted 1/10 for detection of target gene expression. Primer sequences and targets are provided in X. Quantitative real-time PCR was performed with PowerUp SYBR Green Master Mix (Thermofisher, cat. No. A25742). Standard cycling conditions were performed on ViiA 7 Real-Time PCR System according to manufacturer’s instructions. Relative expression of target genes was determined using the ΔΔCt method with *Gapdh* as an internal control.

#### CC-RIX mESC derivation

Collaborative Cross strains were selected on the basis of their genotype across the *Lifr* locus (as wild derived-like [NOD, CAST, PWK, WSB] or classical inbred-like [AJ, B6, 129S1, NZO]) and mESCs were derived using our previously published method (Czechanski et al., 2014) from F1 (CC-RIX) embryos. 20 mESC lines were selected for further analysis as listed in Supplementary Table 4.

#### Self-renewal assays

mESC suspensions were stained with propidium iodide (PI) and flow cytometry was used to isolate and plate 1 cell per well x 2 96 well MEF feeder plates per cell line. After one week of growth, colonies visible under phase contrast microscopy were counted and percent cloning efficiency was estimated as the number of visible colonies divided by the number of cells plated (192).

#### Allele swap

CRISPR/Cas9 donors and guides were designed to engineer reciprocal 129S1 and WSB alleles into the caQTL *Lifr* locus. A 128bp oligo donor was selected, sequences below. A 20bp guide (GGGAGACAGAGCTTTCGACA-129S1 or GGAGACACAGCTTTCGACA-WSB) was cloned into the pSpCas9(BB)-2A-Puro (PX459) V2.0 (Addgene #62988). 2×10^6^ cells seeded on a 60mm dish were co-transfected with equimolar amounts (2.5ug each) of donor and guide vector using Lipofectamine 3000 (Invitrogen #L3000) reagent. After 48 hours, cells underwent puromycin selection and after one week colonies were picked and expanded. Clones were PCR amplified using 5’-AGGCCCAGAGTGATTGACTT-3’ and 5’-GGACACAGCACCCAGATTTC-3’and capillary sequencing of the resulting PCR products was used to identify correct, homozygous targeting.

Donors oligos were as follows: 129S1 SNP, 5’-GAAACACAATAGCAGGCT[T]CAGTTAAGTGACTTTGACCTTGTCGAA-AGCTGTGTCTCCCTTTTGAGATGTTAAAAGTTTCAGGAAAAGAAGCACACTAAAAGAAATCTGGGTGCTGTGTCCACCC-CCA-3’. WSB SNP: 5’-GAAACACAATAGCAGGCT[A]CAGTTAAGTGACTTTGACCTTGTCGAAAGCTCTGTCTCCCTTTTG-AGATGTTAAAAGTTTCAGGAAAAGAAGCACACTAAAAGAAATCTGGGTGCTGTGTCCACCCCCA-3’.

#### LIF dose response

mESCs were thawed onto gelatinized dishes into standard culture media containing Dulbecco’s Modified Eagle Medium (DMEM) supplemented with 15% fetal bovine serum, 100 U/mL Penicillin-Streptomycin, 2mM GlutaMAX, 0.1mM non-essential amino acids, 1mM sodium pyruvate, 0.1mM 2-mercaptoethanol, 500pM LIF, 1uM PD0325901, and 3uM CHIR99021. After 48 hours cells were trypsinized, washed, and counted. 3×10^4^ cells were plated per gelatinized 35mm dish in LIF concentrations ranging from 0.05-5000 pM. A control well containing 480 ng/ml of neutratlizing anti-LIF antibody (R&D Systems AB-449-NA) was also plated. After 5 days cells were dissociated and the expression of mCherry was analyzed using the FACsymphony cytometer. Prior to flow cytometry analysis, the cells were stained with DAPI to determine viability.

## Supplemental Information

### Supplemental Tables

**Table S1, related to Figure 1.** Inbred strain mESC lines and quality control measures, normalized gene expression matrix for founder mESCs, and results of differential expression tests using DESeq2.

**Table S2, related to Figure 2.** Genome coordinates of hotspot QTL and lists of features with QTL peaks that may be regulated by each hotspot.

**Table S3, related to Figure 3.** Candidate mediators for each QTL hotspot.

**Table S4, related to Figure 4.** The Collaborative Cross strains, Chr15 QTL region genotypes, and crosses used to generate CC-RIX mESCs.

### Supplemental Figures

**Figure S1, related to Figure 2.**
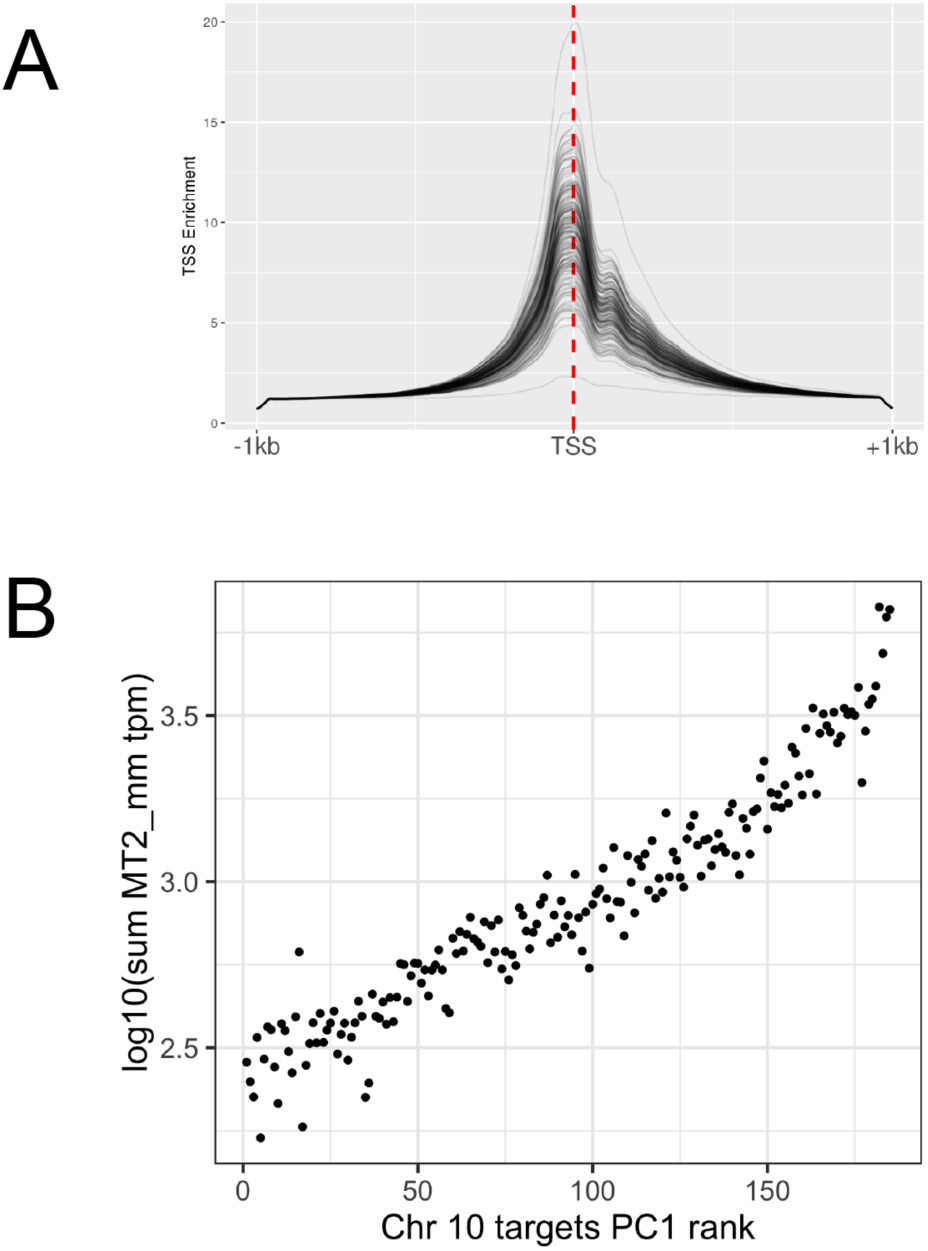
Open chromatin at promoters and transcription of endogenous retroviral elements. (A) Open chromatin at the transcription start site (TSS) across 16,956 genes annotated by Gencode (Frankish et al., 2019). ATAC-Seq read depth was calculated at each TSS +/− 1kb, depth was normalized to one at the -1kb and +1kb boundaries, and relative depth is plotted. Each line shows a separate DO mESC sample. (B) Correlation between transcription of the MERVL long terminal repeat MT2_mm (http://www.dfam.org/entry/DF0004155) and Chr 10 hotspot QTL target genes. Reads from each DO mESC sample were aligned to a transcriptome consisting of all annotated genes (Ensembl release 82) as well all non-redundant mouse genome hits to the Dfam (Hubley et al., 2016) HMM for this repeat element using kallisto (Bray et al., 2016) version 0.43.1. Figure shows samples ranked by principal component 1 of the expression matrix of Chr 10 hotspot QTL targets, plotted against the summed transcripts per million for all MT2_mm transcripts. Individual points represent individual DO mESC samples.

**Figure S2, related to Figure 2.**
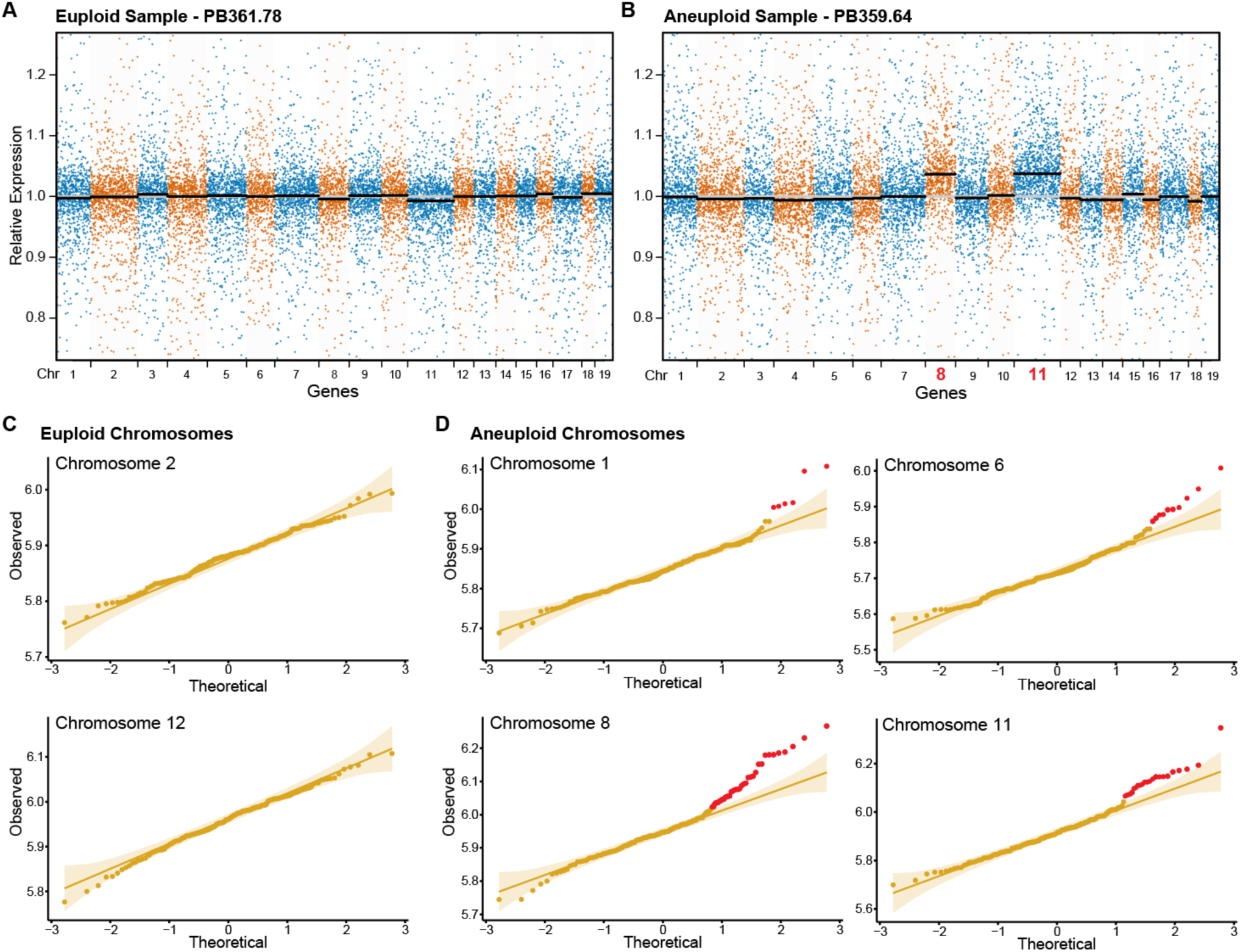
Characterizing the extent of aneuploidy in DO mESC lines. Gene expression data can be used to identify DO lines that have acquired chromosomal duplications. (A-B) Within a sample, each gene’s expression relative to the population median is plotted on the *y*-axis against the gene’s chromosome on the *x*-axis. Within a chromosome, genes are ordered by position (but plotted as equidistant to more clearly display trends). (A) ESC lines that do not have an appreciable proportion of aneuploid cells show the median gene expression of each chromosome nearing the population median (black bars are the sample chromosome medians; grey line is the population median = 1). (B) ESC lines with larger proportions of aneuploid cells deviate from the population median for the duplicated chromosomes (Chromosomes 8 and 11 in panel B). (C-D) The distribution of sample chromosome medians can reveal ESC lines with duplications of a given chromosome. (C) For chromosomes euploid across all samples, we assume that the sample median expression values will fit a normal distribution (representative euploid chromosomes shown in C). (D) Chromosomes that are aneuploid in ESC lines should violate this assumption, with aneuploid samples exhibiting higher median expression values that fall outside the normal distribution. Using these criteria, chromosomes 1, 6, 8, and 11 were classified as aneuploid in multiple DO lines (Shapiro-Wilk Normality Test p > 0.05).

**Figure S3, related to Figure 3.**
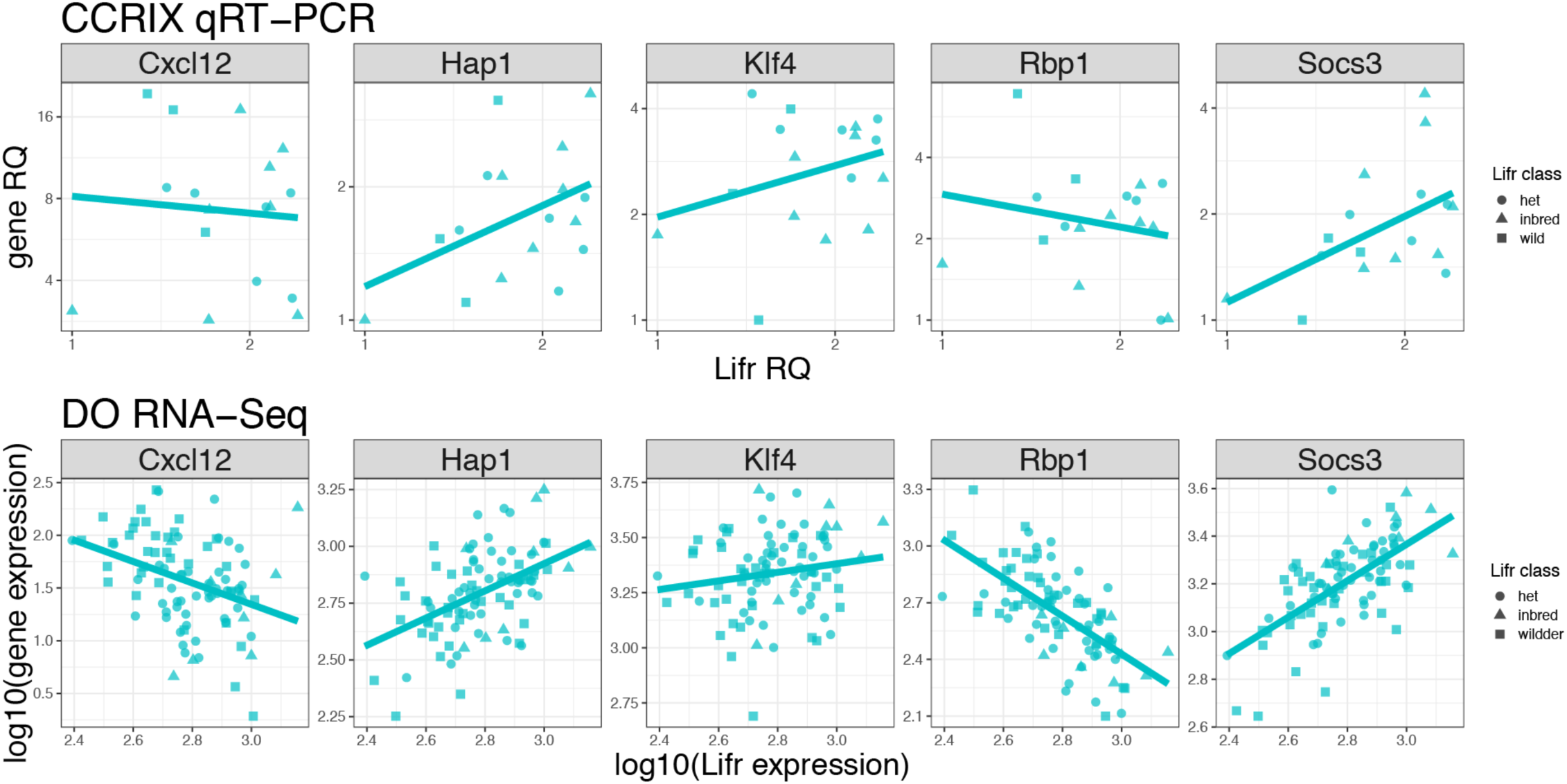
Quantitative PCR validation of five gene expression hotspot targets. qRT-PCR validation of five genes with eQTL mapping to the Chr 15 hotspot. We compared RQ (relative quantification via qPCR) of *Lifr* to each of the other five genes (facets). Top panels show qPCR of CC-RIX mESCs, while bottom panels show gene expression inferred from bulk RNA-Seq of DO mESCs. Only male CC-RIX mESC lines were used, thus data from DO mESC lines is restricted to only male lines. Note that gene co-expression relationships show similar trends (lines show best-fit regression) despite different genetic resource populations and methods of quantitation.

**Figure S4, related to Figure 4.**
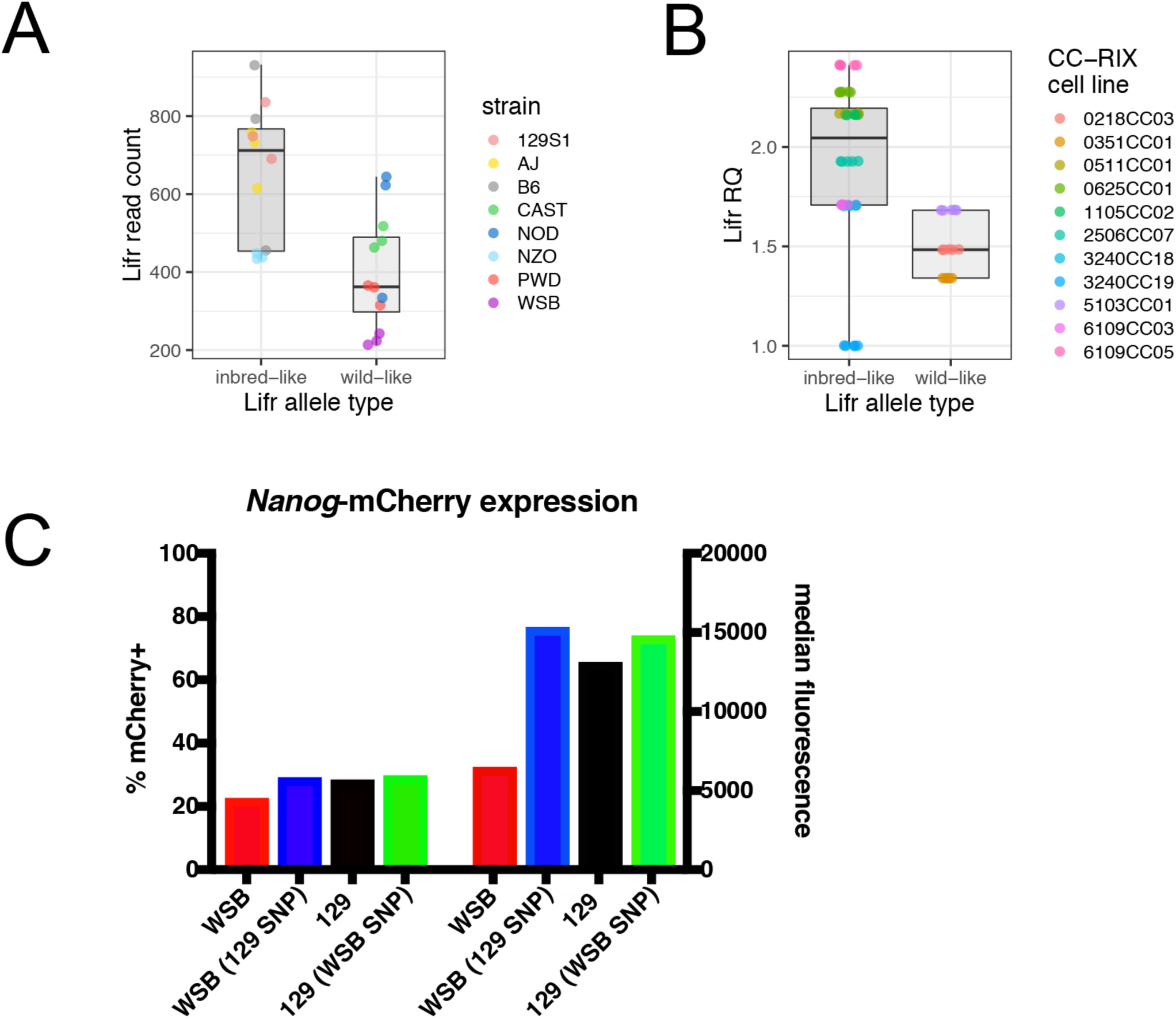
Validating genetically driven variation in *Lifr* expression and the effect of allele swaps on NANOG expression. (A) *Lifr* genotype class (classical inbred-like or wild-derived-like) affects *Lifr* expression as measured by RNA-Seq in CC founder strains. (B) *Lifr* genotype class affects *Lifr* expression as measured by qPCR in CC-RIX mESCs (only homozygous lines are shown). RQ indicates relative quantification via qRT-PCR. (C) Flow cytometry provides quantitative assessment of the percentage of cells expressing and relative intensity of *Nanog*-mCherry expression. Figure shows *Nanog*-mCherry expression in representative 129S1 and WSB parental mESC lines, as well as the allele swap clones. In 2i culture conditions, the WSB parental mCherry lines have a slightly reduced percentage of cells expressing mCherry, but more significantly reduced median fluorescence. The positive effect of the allele swap in WSB cells with regard to the percentage of cells expressing and median fluorescence is apparent, however we do not observe the opposing effect of the allele swap in 129S1.

